# PIMT suppresses endothelial activation and vascular inflammation through methylation of TRAF6

**DOI:** 10.1101/2022.01.25.477656

**Authors:** Chen Zhang, Zhi-Fu Guo, Bing Yi, Kyosuke Kazama, Wennan Liu, Xiaobo Sun, Lu Wang, Xiao-Feng Yang, Ross Summer, Jianxin Sun

**Affiliations:** Center for Translational Medicine, Thomas Jefferson University, Philadelphia, PA, USA; Department of Cell and Developmental Biology, Perelman School of Medicine, University of Pennsylvania, Philadelphia, PA, USA; Center for Metabolic Disease Research, Lewis Katz School of Medicine, Temple University, Philadelphia, PA; Department of Microbiology and Immunology, Lewis Katz School of Medicine, Temple University, Philadelphia, PA

**Author notes:** Correspondence to: Jianxin Sun, Ph.D., Center for Translational Medicine, Thomas Jefferson University, 1020 Locust Street, Room 368G, Philadelphia, PA 19107.

## Abstract

As a repair enzyme, protein L-isoaspartyl O-methyltransferase (PIMT) methylates and converts isoaspartyl residues (isoAsp) back to conventional forms to avoid protein damage during aging and stress responses. This study aims to investigate the pathological significance of PIMT in vascular inflammation. Herein, we show that PIMT is highly expressed in lung endothelial cells (ECs), and its reduction significantly exacerbates pulmonary inflammation in a murine model of acute lung injury (ALI). Mechanistically, we identify tumor necrosis factor receptor-associated factor 6 (TRAF6) as a substrate of PIMT. PIMT-mediated methylation of TRAF6 inhibits TRAF6 oligomerization, autoubiquitination, and LPS-induced NF-κB transactivation in ECs. Importantly, in addition to transcriptionally attenuating expression of adhesion molecules, PIMT post-translationally impedes site specific N-glycosylation of intercellular adhesion molecule-1 (ICAM-1), which leads to an increased protein degradation of ICAM-1 and a subsequent inhibition of EC interaction with immune cells during inflammatory process. Our results for the first time identify PIMT-mediated protein O-methylation as a key post-translational mechanism in controlling vascular inflammation, and suggest that PIMT may represent a novel therapeutic target in inflammatory vascular diseases.

## Introduction

Lung endothelium is a single layer of cells that lines the luminal surface of pulmonary blood vessels and serves an important role in limiting the entry of blood constituents into the lung and controlling growth and inflammatory responses (Galley & Webster, 2004; Pries & Kuebler, 2006). As an early participant in the pathogenesis of acute lung injury and the acute respiratory distress syndrome (ARDS), activation of lung endothelium triggers the expression of pro-inflammatory cytokines and cell surface adhesion molecules to facilitate circulating leukocyte infiltration (Maniatis *et al*, 2008; Millar *et al*, 2016). Unchecked endothelial activation induces excessive inflammatory responses, which ultimately leads to alveolar capillary barrier disruption and lung damage. Interventions targeting endothelial activation and vascular inflammation have emerged as a new paradigm in the treatment of ALI and ARDS (Orfanos *et al*, 2004).

Canonical NF-κB (nuclear factor kappa enhancer binding protein) signaling plays a pivotal role in endothelial cell activation (Baeuerle, 1998). Binding of proinflammatory cytokines to receptors, such as members of interleukin-1 receptor/Toll-like receptor (IL-1R/TLR) superfamily leads to activation of the IKK (IκB [NF-κB inhibitor] kinase) complex via recruiting cytoplasmic adaptors, including TNF Receptor Associated Factors (TRAFs). The activation of the IKK complex causes phosphorylation of IκBα and its polyubiquitination and subsequent proteasomal degradation. The liberated NF-κB dimer then translocates to the nucleus wherein it binds to specific DNA sequences to promote expression of cytokines and endothelial adhesion molecules (Dunne & O’Neill, 2003; Szmitko *et al*, 2003). TRAF6 is a RING-domain cognate ubiquitin ligase (E3) associated with the intracellular domain of the interleukin-1 receptor and Toll-like receptor (IL-1R/TLR) and mediates immune responses through lysine 63 (K63)-linked ubiquitination of diverse signaling intermediators, including itself (Chen, 2005; Deng *et al*, 2000). Negative regulators of TRAF6 have been shown to suppress cellular inflammation, suggesting clinical potential in the treatment of inflammatory diseases (Lv *et al*, 2018; Wang *et al*, 2006).

As another hallmark of the vascular inflammatory process, leukocyte-endothelial adhesion is mediated by interactions between endothelial adhesion molecules and their cognate receptors on immune cells (Szmitko *et al*., 2003). Intercellular adhesion molecule-1 (ICAM-1; CD54) is an inducible transmembrane protein expressed on the surface of endothelial cells, with known roles in leukocyte adhesion and transmigration (Springer, 1994; Staunton *et al*, 1988). Besides, ligand binding conferred ICAM-1 cross-linking functions as the signal transducer to promote proinflammatory effects (Lawson & Wolf, 2009). ICAM-1 is heavily N-linked glycosylated on its extracellular Ig-like domains, and previous studies have demonstrated that this type of post-translational modification (PTM) is indispensable for the proper conformation and biological function of ICAM-1 during inflammatory responses (Chen *et al*, 2014b). However, the molecular mechanisms underlying the regulation of ICAM-1 glycosylation remains largely unknown.

As a conservative protein repair enzyme, protein L-isoaspartyl O-methyltransferase (PIMT) utilizes S-Adenosyl methionine (SAM/AdoMet) to methylate covalently modified isoaspartate (isoAsp) residues accumulated in proteins via Asn deamidation and Asp hydrolysis under physiological or stressed conditions. Apart from normal cell physiology, formation of isoAsp in proteins is promoted by various stress conditions and usually disturbs protein structure and function (Geiger & Clarke, 1987; Reissner & Aswad, 2003), thus conversion of isoAsp to Asp by PIMT plays an essential role in restoring protein function (Desrosiers & Fanélus, 2011; Mishra & Mahawar, 2019; Reissner & Aswad, 2003). It is increasingly recognized that asparagine deamidation is significantly engaged in diverse biological functions such as cell apoptosis and the control of metabolic responses under inflammatory and stress conditions (Deverman *et al*, 2002; Zhao *et al*, 2020), PIMT-mediated protein carboxyl methylation (*O*-methylation) was found to be prevailing in cells, indicating it may functionally rival other types of protein PTMs (Sprung *et al*, 2008). For instance, overexpression of PIMT has been shown to extend longevity of bacteria, fruit flies and plants (Chavous *et al*, 2001; Ogé *et al*, 2008; Pesingi *et al*, 2017). PIMT deficient mice exhibit increased isoAsp deposition and early death induced by grand mal seizures (Kim *et al*, 1997). Furthermore, we and others have shown that PIMT plays an important role in the regulation of angiogenesis and cardiomyocyte survival under oxidative conditions (Ouanouki & Desrosiers, 2016; Yan *et al*, 2013).

Here, we demonstrate that PIMT is a potent negative regulator of endothelial activation and vascular inflammation. Mechanistically, we identify TRAF6 as a novel substrate of PIMT and PIMT-mediated methylation of TRAF6 markedly inhibits LPS-induced NF-κB activation in ECs. Furthermore, we found that PIMT interacts with ICAM-1, which in turn inhibits ICAM-1 N-glycosylation and its functional interaction with immune cells. Our study provides a novel mechanism for controlling EC activation at posttranslational levels through PIMT-mediated protein *O*-methylation.

## Results

### 1. PIMT mitigates LPS-induced mouse pulmonary vascular inflammation and lung injury

Lung infection sustained by various pathogens, such as bacteria and the coronavirus (COVID-19) is the leading cause of ALI/ARDS, and its severity and morbidity are markedly increased in the elderly (Akbar & Gilroy, 2020; Kang & Jung, 2020). Despite its function as an anti-aging and stress protein, the role of PIMT in ALI/ARDS is still unknown. In this regard, we sought to investigate the pathological significance of PIMT in a murine model of ALI induced by LPS instillation. Since homozygous deletion of both PIMT alleles leads to fatal epileptic seizures at 30-60 days after birth (Kim *et al*., 1997), PIMT^+/-^ mice and their WT counterparts were deployed in this study. Following LPS instillation, we found that the susceptibility of mice to ALI is significantly increased in PIMT^+/-^ mice. This correlated with increased total protein levels and numbers of leukocytes and neutrophil cells in isolated bronchoalveolar lavage fluid (BALF) from PIMT^+/-^ mice compared with their WT counterparts (Figure 1A and 1B). Furthermore, we found that levels of BALF cytokine and chemokine of tumor necrosis factor alpha (TNF-α), interleukin-6 (IL-6) and C-C motif chemokine ligand 2 (Ccl2), as detected by enzyme-linked immunosorbent assay (ELISA), were also significantly increased in PIMT^+/-^ mice compared with that of WT mice (Figure 1C). To further evaluate the role of PIMT in pulmonary inflammation, we examined the expression of adhesion molecules and inflammatory cytokines in lung tissues. Baseline expression of TNF-α, interleukin 1 beta (IL-1β), IL-6, and Ccl2 was indistinguishable between PIMT^+/-^ and WT mice. However, after LPS instillation, their mRNA levels were significantly increased in PIMT^+/-^ mice compared with the WT controls (Figure 1D). Likewise, the expression of adhesion molecule ICAM-1 was significantly increased in PIMT^+/-^ mice compared with WT counterparts, as determined by both qRT-PCR and western blot (Figure 1E). Histological analysis demonstrated exaggerated lung injury in PIMT^+/-^ mice, as manifested by the presence of increased alveolar hemorrhage, infiltration of inflammatory cells into distal air sacs, and disruption of alveolar walls in response to LPS stimulation (Figure 1F). Importantly, immunofluorescent staining indicate that PIMT is highly expressed in pulmonary vascular endothelium (Figure 1G), suggesting a potential role of endothelial PIMT in regulating lung inflammation. Together, these findings suggest that PIMT prevents inflammatory lung damage potentially through inhibiting pulmonary endothelial activation.

**Figure 1.**
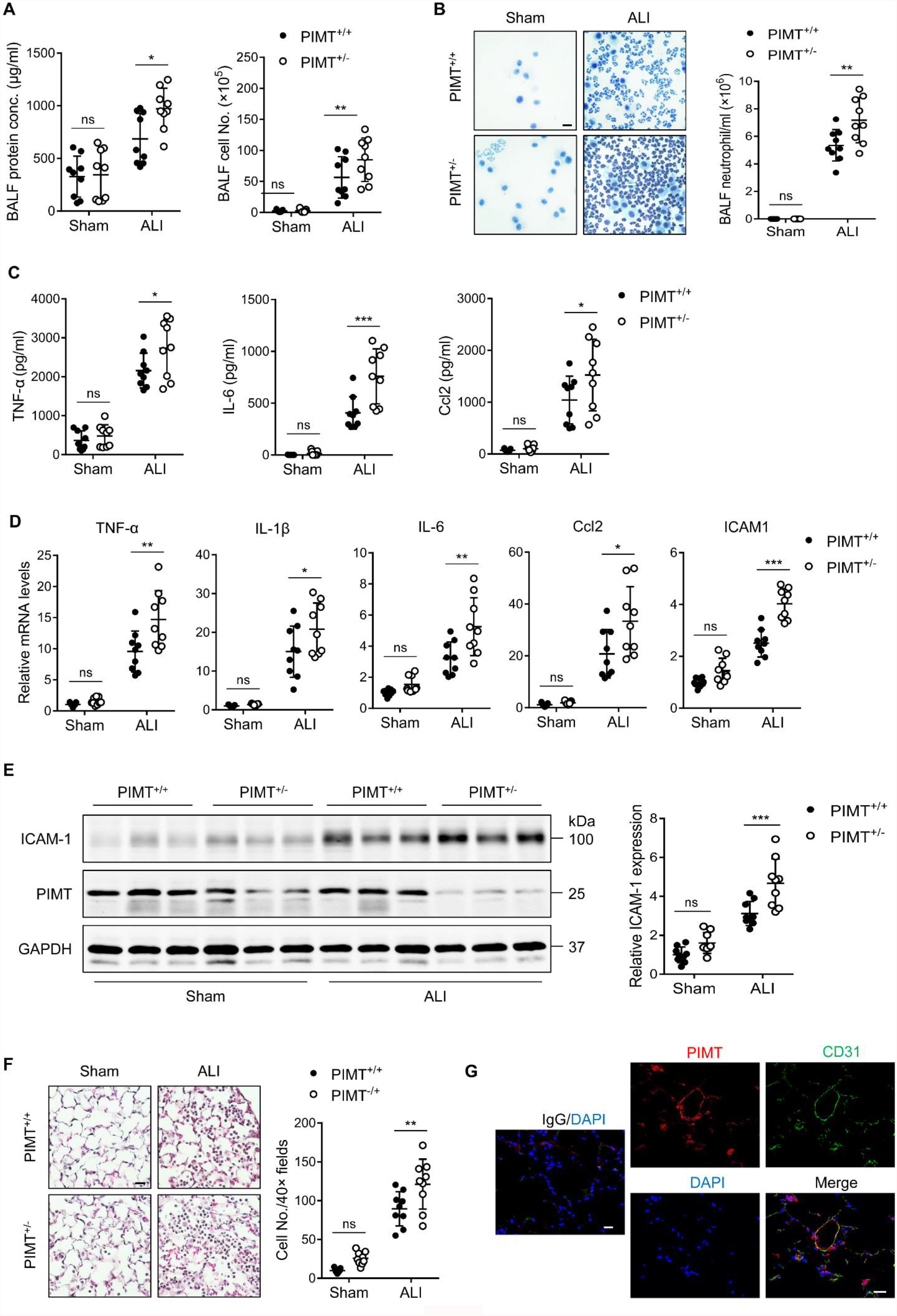
PIMT haploinsufficiency exacerbates LPS-induced mouse pulmonary vascular inflammation and lung injury. **A**, Total cell counts and protein concentrations were determined in BALF of WT (PIMT^+/+^) and PIMT hemizygous (PIMT^+/-^) mice 18 hrs after intratracheal LPS (100 μg/100 μl, ALI) or PBS (100 μl, Sham) administration. **B**, Diff-Quik stained smears of BALF (left) and neutrophil counts (right) were determined in WT and PIMT^+/-^ mice 18 hrs after intratracheal PBS or LPS administration. Bars, 20 μm. **C**, Levels of TNF-α, IL-6 and Ccl2 in the BALF of WT and PIMT^+/-^ mice were measured by ELISA 18 hrs after intratracheal PBS or LPS administration. **D**, Expression of TNF-α, IL-1β, IL-6, Ccl2 and ICAM-1 mRNA extracted from lung tissues of WT and PIMT^+/-^ mice with indicted treatments were determined by qRT-PCR. **E**, Expression of ICAM-1 from lung tissues of WT and PIMT^+/-^ mice with indicted treatments were determined, shown by representative western blot (left) with the quantitative results (right). **F**, Hematoxylin and eosin (H&E) staining of the lung sections of WT and PIMT^+/-^ mice 18 hrs after intratracheal PBS or LPS treatment. Infiltrated immune cells were counted. Bars, 20 μm. **G**, Representative immunofluorescent staining of PIMT in WT mouse lung sections. CD31 is shown an endothelial marker. Nuclei were stained with DAPI (4′,6-diamidino-2-phenylindole), IgG was used as a negative control. Bars, 20 μm. All data are expressed as mean ± SD, \**P* < 0.05, \*\**P* < 0.01, \*\*\**P* < 0.001, two-way ANOVA coupled with Tukey’s post hoc test, n = 9 per group.

### 2. PIMT inhibits endothelial NF-κB transactivation

LPS has been known to stimulate the expression of inflammatory molecules in ECs via the TLR-NF-κB pathway (Dauphinee & Karsan, 2006). To define whether PIMT plays a role in regulating endothelial activation, we examined the effect of PIMT overexpression on NF-κB promoter activity at the transcriptional level (Ahn *et al*, 1995). EA.hy926 endothelial cells were transiently transfected with a plasmid containing a heterologous promoter driven by NF-κB elements upstream of the luciferase gene. As shown in Figure 2A and B, overexpression of PIMT markedly reduced the NF-κB responsive under both basal and LPS-stimulated conditions. By contrast, ectopic expression of an inactive PIMT mutant, which has a deletion in the catalytic motif I (PIMT-Del), barely affected LPS-induced NF-κB activation, suggesting that the PIMT enzymatic activity is indispensable for this inhibitory function.

**Figure 2.**
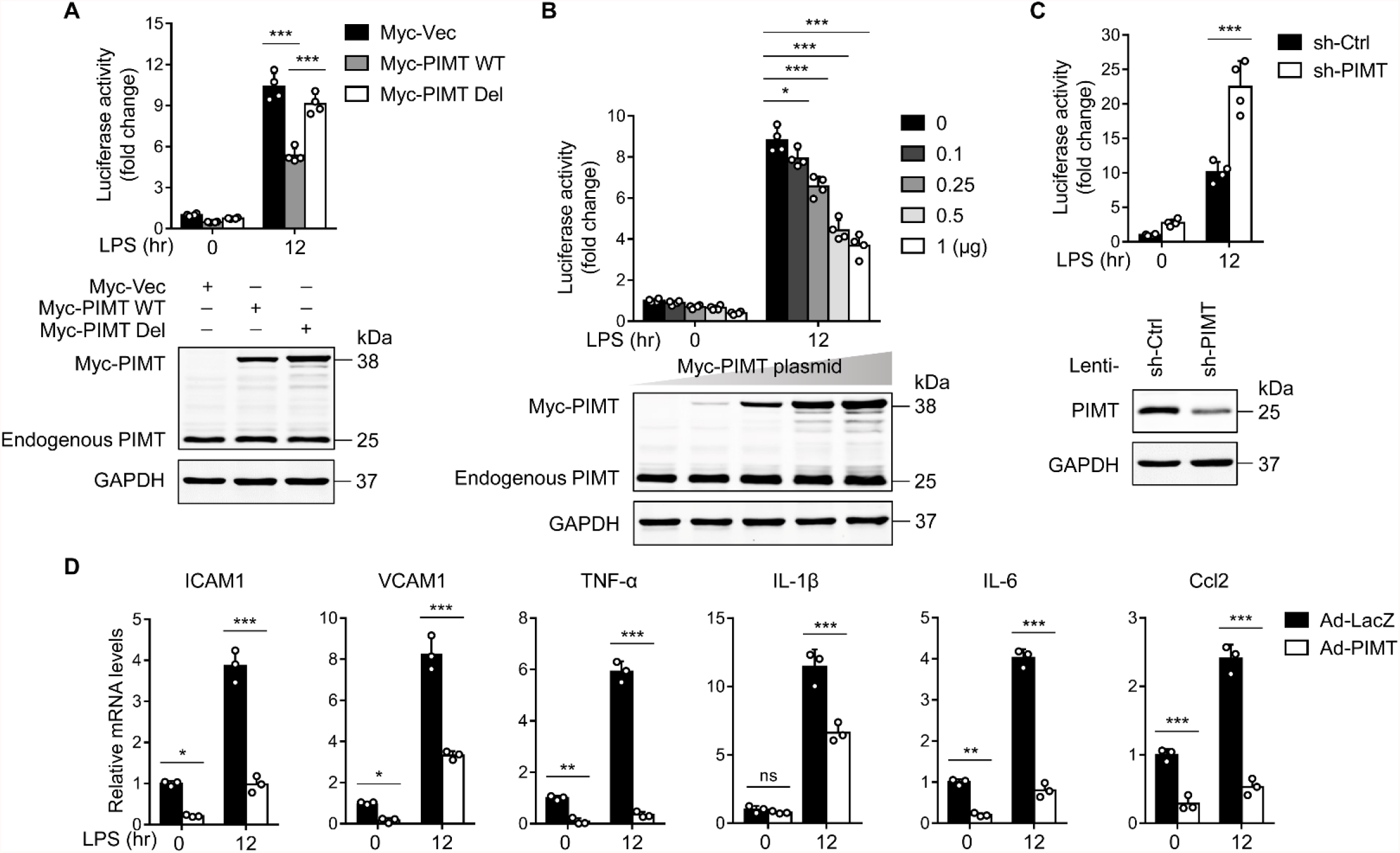
PIMT suppresses LPS-induced endothelial NF-κB transactivation and pro-inflammatory effect. **A**, EA.hy926 cells were transfected with NF-κB reporter together with empty or Myc-tagged PIMT plasmids. 36 hrs after transfection, cells were treated with either vehicle or LPS (1 μg/ml) for additional 12 hrs, then harvested for the luciferase assay. Results are presented relative to Renilla activity (n = 4). Endogenous and ectopic expression of PIMT were measured by western blot. Vec, pCS2-6×Myc empty vector; WT, wild-type PIMT; Del, catalytic defective PIMT mutant. **B**, EA.hy926 cells were co-transfected with NF-κB reporter together with increased amount of Myc-tagged PIMT plasmids. 36 hours after transfection, cells were stimulated with either vehicle or LPS (1 μg/ml) for 12 hours and then harvested for luciferase assay (n = 4). Expression of PIMT was determined by western blot. **C**, PIMT stable depleted (sh-PIMT) and non-targeting control (sh-Ctrl) EA.hy926 cells were transfected with NF-κB reporter plasmid for 36 hrs, luciferase activity was determined after LPS (1 μg/ml) treatment for 12 hrs (n = 4). PIMT levels were detected by western blot. **D**, HUVECs were transduced with adenovirus LacZ (Ad-LacZ) or PIMT (Ad-PIMT) at a multiplicity of infection (moi) of 10 for 48 hrs. The mRNA expression of ICAM1, VCAM1, TNF-α, IL-1β, IL-6 and Ccl2 were determined by qRT-PCR after LPS (1 μg/ml) treatment for 12 hrs (n = 3). Data are representative of mean ± SD, \**P* < 0.05, \*\**P* < 0.01, \*\*\**P* < 0.001, two-way ANOVA coupled with Tukey’s post hoc test.

Similarly, the inhibitory effects of PIMT on NF-κB activation were observed in TNF-α- or IL-1β-stimulated cells (Supplementary Figure 1A and B). To further elucidate the role of PIMT in NF-κB activation, we performed loss of function studies by using a lentivirus expressing PIMT shRNA (sh-PIMT) or non-targeting control shRNAs (sh-Ctrl). As shown in Figure 2C, knockdown of PIMT expression by lentiviral sh-PIMT significantly augmented LPS-stimulated NF-κB promoter-driven luciferase activity. Similar effects were also observed in IL-1β-treated HEK-293T cells transfected with PIMT small interfering RNA (siRNA) (Supplementary Figure 1C). Furthermore, we determined the effects of PIMT on the expression of adhesion molecules, cytokines, and chemokines in ECs. As shown in Figure 2D, adenovirus-mediated overexpression of PIMT markedly attenuated the expression of ICAM1, vascular cell adhesion molecule 1 (VCAM1), IL-1β, IL-6, IL-8, and Ccl2 under both basal and LPS-stimulated conditions, as determined by qPCR. Collectively, these results suggest that PIMT is a novel negative regulator of NF-κB activation in ECs.

### 3. PIMT attenuates endothelial TLR signaling

Canonical NF-κB pathway is transduced from cell surface receptors to nuclear events, and various effectors are implicated for this cascade signaling (Lawrence, 2009). To further identify the components in NF-κB signaling pathway regulated by PIMT, we examined whether PIMT impacts phosphorylation of IKKβ and IκBα, two important components implicated in NF-κB activation. As shown in Figure 3A, PIMT overexpression significantly attenuated phosphorylation of IKKβ and IκBα at different time points after LPS stimulation, compared with ECs transduced with control adenovirus (Ad-LacZ). Furthermore, we found that PIMT depletion led to a marked increase of phosphorylation of IKKβ and IκBα in ECs compared with lentiviral expressing control shRNA (Figure 3B). Moreover, LPS-induced nuclear translocation of the p65 subunit of NF-κB, as determined by western blot (Figure 3C) and immuofluorescent staining, was significantly reduced in PIMT overexpressing cells (Figure 3D). Accordingly, NF-κB DNA binding capacity, as determined by electrophoretic mobility-shift assay (EMSA), was significantly inhibited by ectopic expression of PIMT in ECs (Figure 3E). Together, these results suggest that PIMT inhibits LPS-induced endothelial NF-κB activation through acting on IKKβ kinase and/or its upstream regulators.

**Figure 3.**
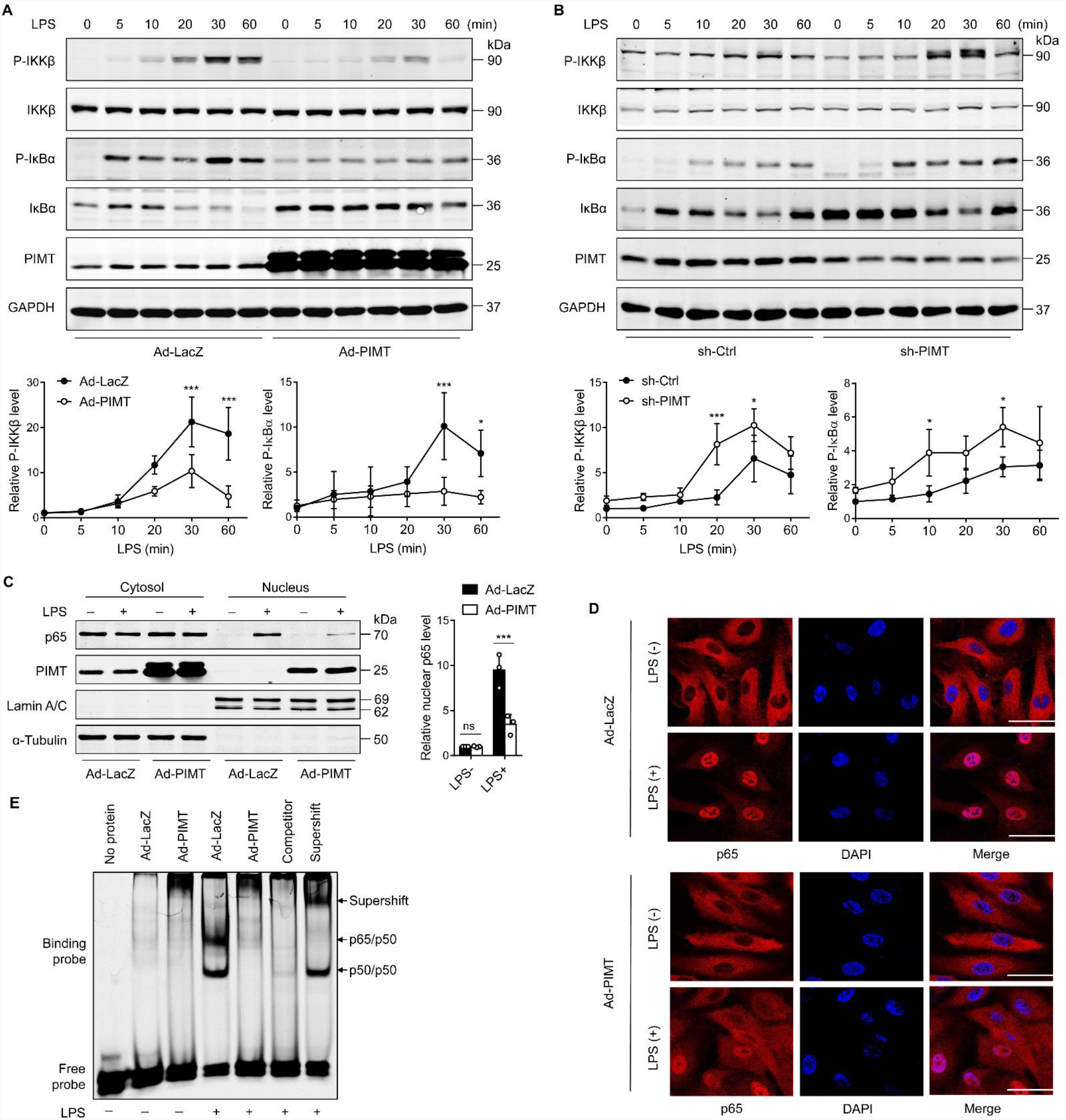
PIMT attenuates endothelial TLR signaling. **A**, HUVECs were transduced with Ad-LacZ or Ad-PIMT (moi = 10) for 48 hrs and then stimulated with LPS (1 μg/ml) for indicated time points. Expression of PIMT and phosphorylation of the indicated proteins in TLR-mediated NF-κB pathway were detected by western blot (above) and quantitated by densitometric analysis (below). **B**, HUVECs were transduced with lentiviral control (sh-Ctrl) or PIMT shRNA (sh-PIMT) for 72 hrs and stimulated with LPS (1 μg/ml) for indicated time points. Expression and phosphorylation of indicated proteins were determined by western blot (above) and quantitated by densitometric analysis (below). **C**, HUVECs were transduced with Ad-LacZ or Ad-PIMT (moi=10) for 48 hrs, and then treated with LPS (1 μg/ml) for 30 mins. Levels of NF-κB p65 and p50/p105 in the cytoplasmic and nuclear fractions were determined by western blot. Lamin A/C and α-Tubulin were used as nuclear and cytoplasmic markers. The representative images (left) and quantification (right) were shown. **D**, HUVECs were transduced with Ad-LacZ or Ad-PIMT (moi = 10) for 48 hrs, followed by LPS (1 μg/ml) stimulation for 30 mins. Localization of p65 was determined by immunofluorescent staining. Nuclei were stained with DAPI. Scale bars, 20 μm. **E**, HUVECs were transduced with Ad-LacZ or Ad-PIMT (moi = 10) for 48 hrs and then stimulated with LPS (1 μg/ml) for 30 mins. Nuclear fraction was extracted and NF-κB DNA-binding activity was determined by EMSA. Competitor was 50-fold unlabeled probe. Supershift was incubated with anti-p65 antibody. All data are representative of mean ± SD, \**P* < 0.05, \*\*\**P* < 0.001, two-way ANOVA coupled with Tukey’s post hoc test, n=3.

### 4. PIMT interacts with TRAF6

TRAFs are critically involved in TLR/IL-1 and TNF receptors (TNFRs) signaling pathways that lead to NF-κB activation (Chen, 2005). To define the regulatory targets of PIMT, we performed a luciferase assay by co-transfection of NF-κB reporter plasmid together with expression vectors of TRAF2, TRAF6, and IKKβ. As shown in Figure 4A, ectopic expression of PIMT significantly inhibited the NF-κB reporter activation induced by TRAF2 and TRAF6, but not by IKKβ-SS/EE, a constitutively active mutant of IKKβ (Nottingham *et al*, 2014) in ECs. Furthermore, we examined the interaction of PIMT with TRAF2, TRAF6, and IKKβ by immunoprecipitation using cell lysates from HEK-293T cells exogenously expressing tagged proteins. Our co-immunoprecipitation (co-IP) demonstrated that PIMT was substantially enriched in the anti-TRAF6 immunocomplex as compared with anti-TRAF2 or anti-IKKβ immunocomplexes (Figure 4B). The binding ability of PIMT with TRAF6 was further confirmed by forward and reciprocal co-IP assays (Figure 4C and D). The endogenous interaction of TRAF6 with PIMT was confirmed by co-IP in ECs (Figure 4E and F). Importantly, this endogenous interaction was augmented by LPS stimulation in both HUVECs and human lung microvascular endothelial cells (HLMVECs) (Figure 4F and G), indicating that the dynamic interaction of PIMT and TRAF6 may be functionally important for LPS-induced inflammatory signaling in ECs.

**Figure 4.**
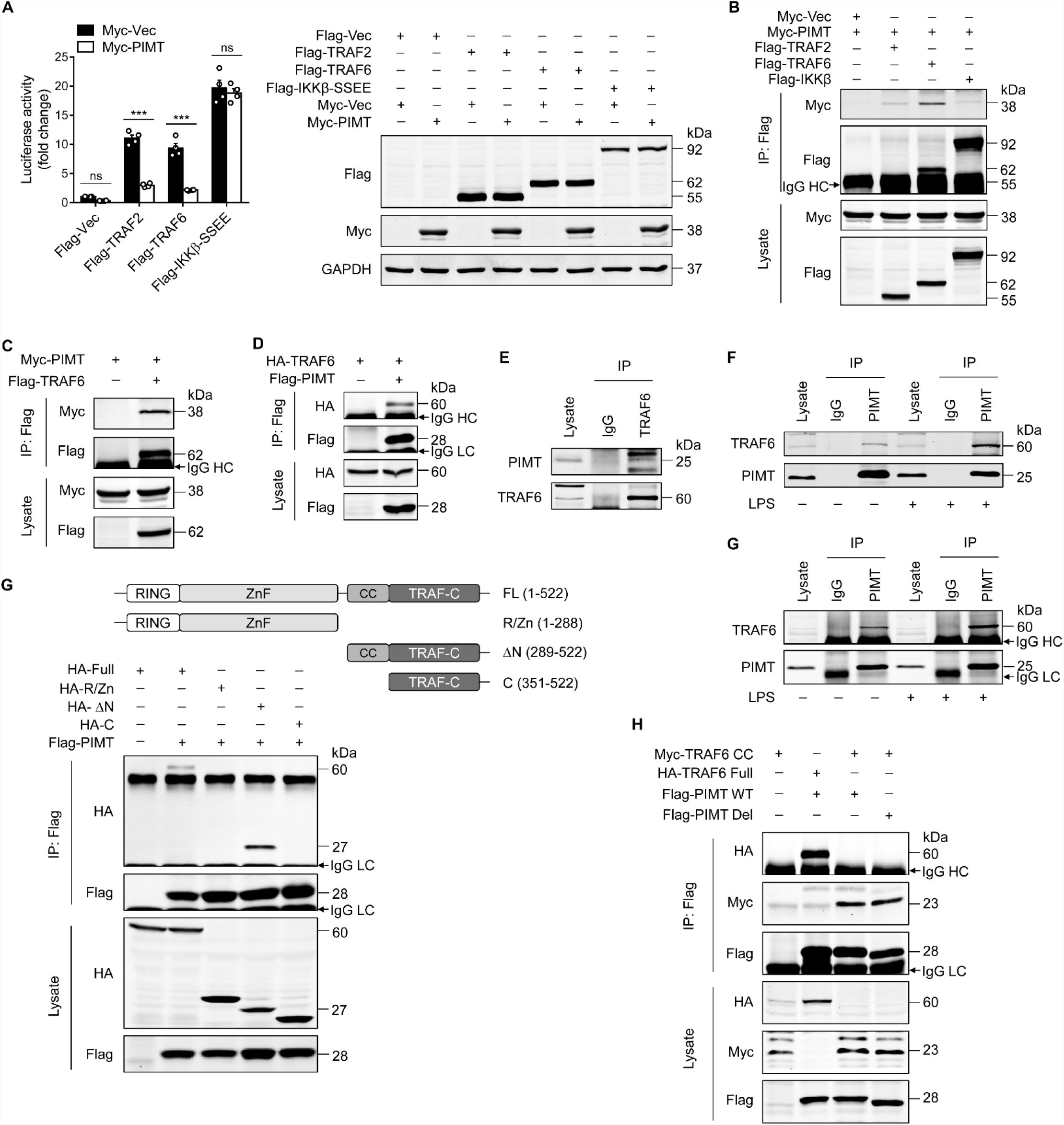
PIMT functionally interacts with TRAF6. **A**, EA.hy926 cells were transfected with NF-κB reporter together with indicated plasmids. Cells were harvested 48 hrs later to determine luciferase activity (n = 4), \*\*\**P* < 0.001, two-way ANOVA coupled with Tukey’s post hoc test. Expression of transfected proteins was detected by western blot. **B**, HEK-293T cell were transfected with a combination of indicated plasmids. 48 hrs after transfection, immunoprecipitation was performed using anti-Flag antibody, followed by western blot to detect protein interaction. **C**, HEK-293T cells were transfected with Flag-TRAF6 and Myc-PIMT expression vectors. Immunoprecipitation was performed 48 hrs later using anti-Flag antibody, followed by western blot to detect the interaction of PIMT with TRAF6. **D**, HEK-293T cells were transfected with HA-TRAF6 and Flag-PIMT vectors. 48 hrs after transfection, Immunoprecipitation was performed using anti-HA antibody, followed by western blot to detect protein interaction. **E**, Immunoprecipitation of HUVEC lysate using anti-TRAF6 antibody, the interaction of TRAF6 with PIMT was determined by western blot. **F**, HUVECs were stimulated with vehicle or LPS (1 μg/ml) for 30 mins. Cell lysates were then collected for immunoprecipitation with anti-PIMT antibody, followed by western blot to detect the interaction of target proteins. **G**, Immunoprecipitation of PIMT and western blot of TRAF6 in cell lysates from HLMVECs with or without LPS (1 μg/ml) treatment for 30 mins. IgG heavy chain (HC) and IgG light chain (LC) were indicated. **H**, Above, Schematic representation of full length (FL) and truncated mutants of TRAF6. Below, HEK-293T cells were transfected with Flag-PIMT together with HA-tagged full length TRAF6 or its deletion mutants. 48 hrs after transfection, immunoprecipitation was performed using anti-Flag antibody, followed by western blot to detect the interaction of PIMT with WT or truncated TRAF6. **I**, HEK-293T cell were transfected with PIMT or its mutants together with tagged TRAF6 or its CC domain. Immunoprecipitation was performed using anti-Flag antibody, followed by western blot to detect the interaction of indicated proteins.

TRAF6 is an ubiquitin E3 ligase which consists of a RING domain at its N-terminus, followed by zinc finger (ZnF) domains, a coiled-coil (CC) domain and a TRAF-C domain at the C-terminus (Chung *et al*, 2002). To map the interacting domain(s) of PIMT in TRAF6, we constructed TRAF6 deletion mutants and determined their interactions with PIMT by immunoprecipitation. In this regard, HEK-293T cells were co-transfected with HA-tagged full-length/truncated TRAF6 and Flag-tagged PIMT plasmids, and the interaction of PIMT with TRAF6 mutants was determined by co-IP using anti-Flag antibody. As shown in Figure 4H, TRAF6 mutant lacking the N-terminal RING domain and ZnF domains was efficiently pulled down by PIMT, while further deletion of the CC domain completely abolished the interaction of PIMT with TRAF6, indicating that the CC domain is essentially involved in the interaction of PIMT with TRAF6. This observation was further confirmed in HEK-293T cells overexpressing Flag-tagged PIMT and Myc-tagged CC domain by immunoprecipitation (Figure 4I). Together, our results demonstrate that the CC domain of TRAF6 is responsible for the binding of PIMT in TRAF6.

### 5. PIMT negatively regulates TRAF6 function through methylation of asparagine at position 350

The CC domain of TRAF6 has been shown to mediate TRAF6 oligomerization and its interaction with ubiquitin-conjugating enzymes, which is essential for the subsequent auto-ubiquitination leading to NF-κB activation (Wang *et al*, 2001; Wang *et al*., 2006). Thus, the interaction of PIMT with TRAF6 CC domain prompted us to test whether PIMT affects TRAF6 oligomerization and auto-ubiquitination. As shown in Figure 5A, Flag-TRAF6 precipitated HA-TRAF6 in co-transfected HEK-293T cells, however, this interaction was markedly inhibited by wild-type PIMT (PIMT-WT), but not by inactive PIMT mutant (PIMT-Del). Similarly, the polyubiquitination of TRAF6 was substantially inhibited by PIMT-WT, but not by PIMT-Del (Figure 5B), suggesting that the enzymatic activity of PIMT is required for the inhibition of TRAF6 activation.

**Figure 5.**
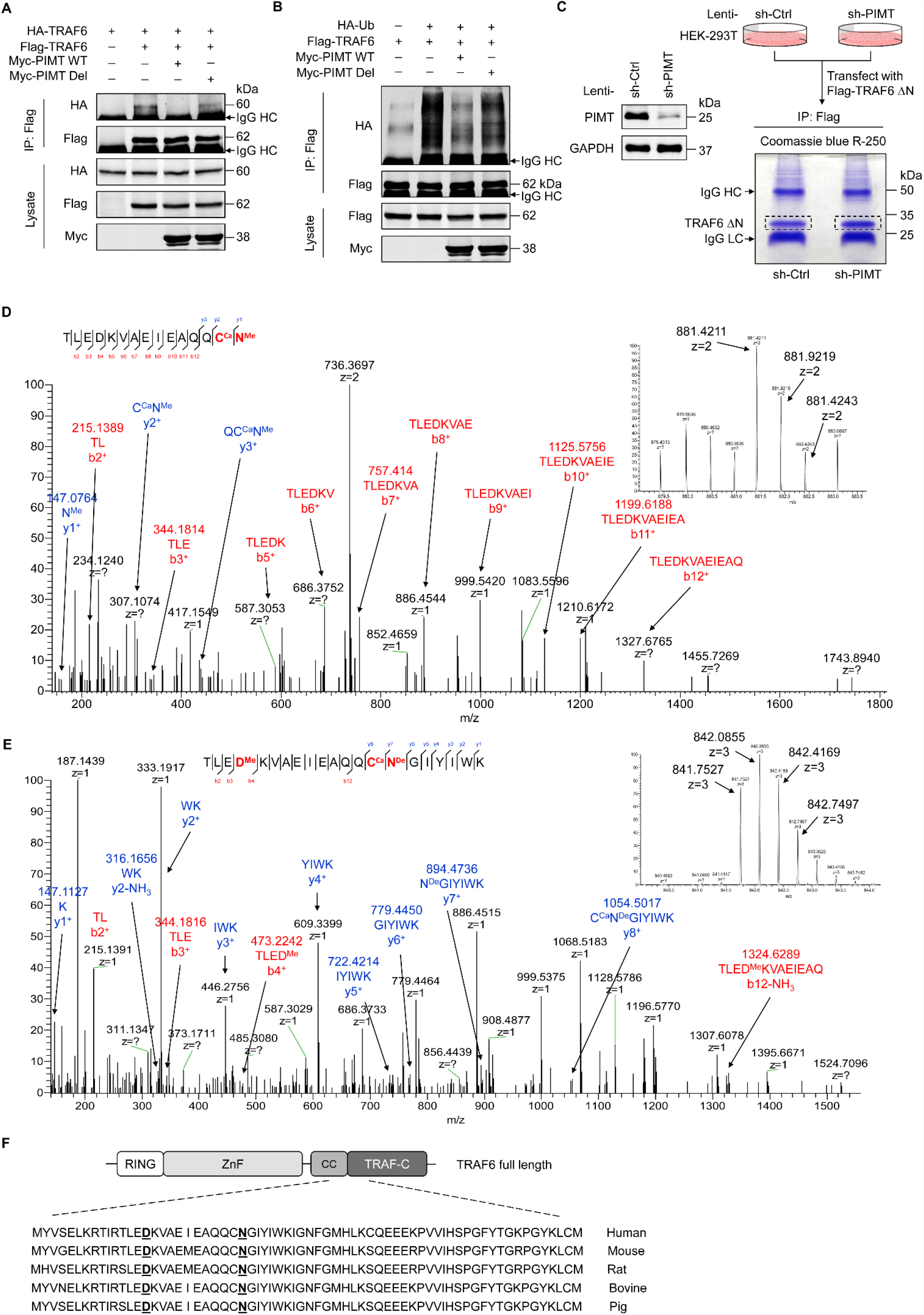
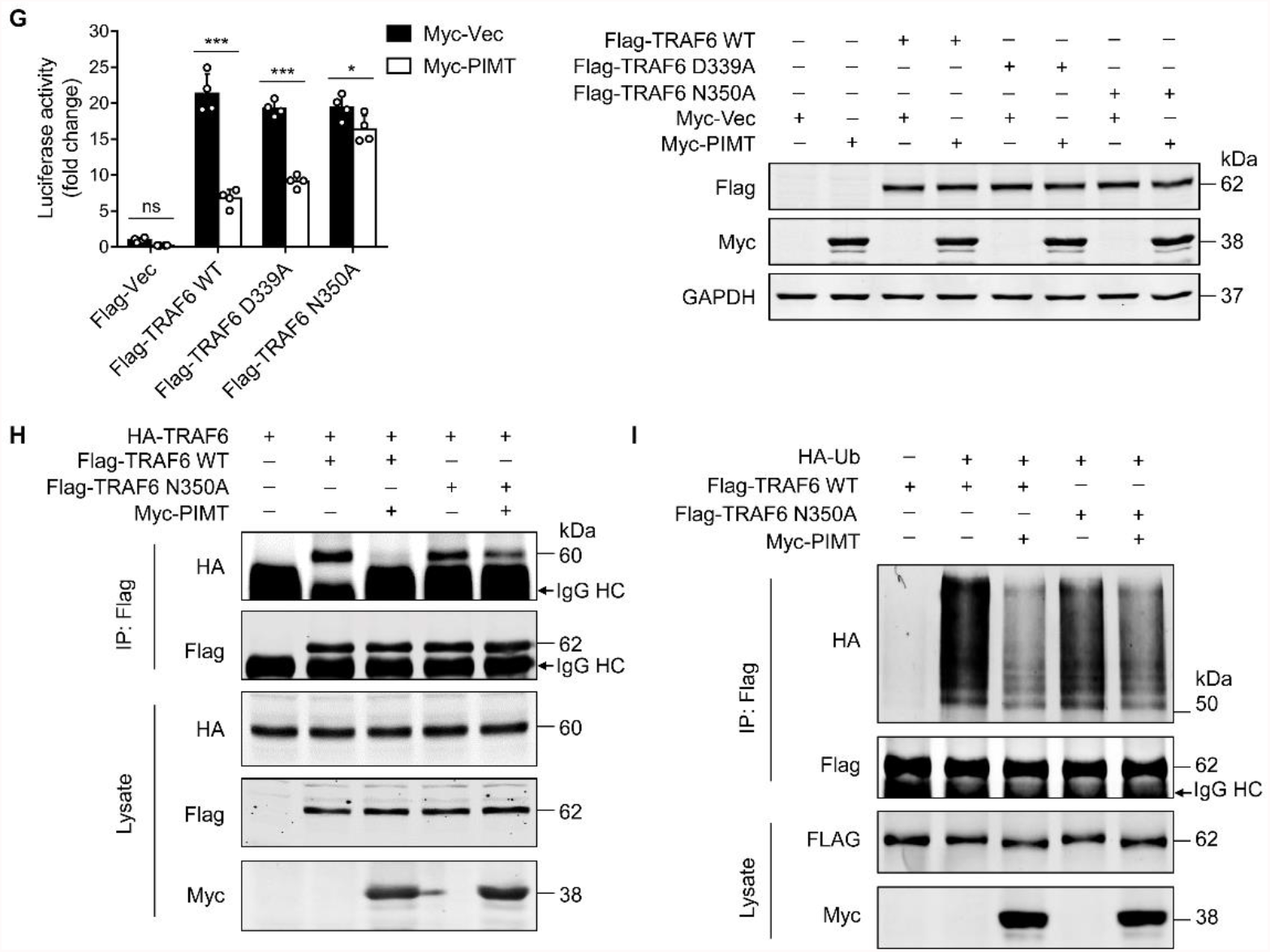
PIMT suppresses TRAF6 function through methylation of TRAF6 CC domain at N350. **A**, HEK-293T cells were transfected with TRAF6 and PIMT constructs. Immunoprecipitation was performed 48 hrs later using anti-Flag antibody, followed by western blot using anti-HA antibody to detect TRAF6 oligomerization. **B**, HEK-293T cells were transfected with HA-ubiquitin (HA-Ub) and Flag-TRAF6 in the presence of Myc-tagged wild type (WT) or mutant PIMT (PIMT Del). 48 hrs after transfection, cells were collected for for anti-Flag immunoprecipitation. The autoubiquitination of TRAF6 was detected by western blot using anti-HA antibody. **C**, Left, western blot showing expression of PIMT in control or PIMT stable depleted HEK-293T cells. Right, work flow of in gel sample preparation of TRAF6 ΔN (shown in Figure 4G) for mass spectrometric analysis. **D** and **E**, LC-MS/MS analysis of TARF6 ΔN domain. Mass tolerance window of the fragment ions is set within 0.02 Da. Ca, carbamidomethyl. Me, methyl. De, deamidated. **D**, (left) manually assigned MS/MS spectra of TLEDKVAEIEAQQC (carbamidomethyl) N (methyl). (Right) corresponding 2 charged precursor ion peaks, the m/z of the monoisotopic peak (m/z 841.4211, z = 2) was found to be 1.4 ppm apart from the theoretical m/z (881.4198, z=2) **E**, (Left) manually assigned MS/MS spectra of TLED (methyl) KVAEIEAQQC (carbamidomethyl) N (deamidated) GIYWK. (Right) corresponding 3 charged precursor ion peaks, the m/z of the monoisotopic peak was exactly matched to the theoretical m/z (841.7527, z = 3). All modified sites were shown in red font. **F**, Alignment of TRAF6 protein sequence from different species. Identified methylated Asp and Asn residues are underlined. **G**, EA.hy926 cells were transfected with NF-κB reporter plasmid together with wild-type TRAF6 (WT) or TRAF6 mutants (D339A and N350A). 48 hrs after transfection, cell were collected for luciferase assay (n = 4), \**P* < 0.05, \*\*\**P* < 0.001, two-way ANOVA coupled with Tukey’s post hoc test. Expression of transfected proteins were determined by western blot. **H**, HEK-293T cells were transfected with HA-TRAF6 and indicated Flag-tagged TRAF6 constructs together Myc-PIMT plasmid. Immunoprecipitation was performed 48 hrs later using anti-Flag antibody, followed by western blot using anti-HA antibody to detect TRAF6 oligomerization. **i**, HEK-293T cells were transfected with Flag tagged WT or mutant TRAF6 with HA-Ub and Myc-PIMT plasmids. 48 hrs after transfection, TRAF6 was immunoprecipitated by anti-Flag antibody and autoubiquitination was then detected by western blot using anti-HA antibody.

The amino acid aspartate (Asp, D) or asparagine (Asn, N) following by small or flexible side chain amino acids such as glycine, histidine or serine residing on protein surface structures are inclined to undergo deamidation yielding isoAsp, which could be methylated by PIMT (Geiger & Clarke, 1987; Reissner & Aswad, 2003). To determine whether PIMT methylates TRAF6 under physiological conditions, TRAF6-ΔN was immunoprecipated from either PIMT expressing or PIMT depleting HEK-293T cells using lentivirus PIMT shRNA (Figure 5C, right), and then subjected to Nano liquid chromatography and mass spectrometry (nano LC-MS/MS) analysis to identify potential methylation sites. Our results demonstrated that D300, D339 and N350, located in the CC domain (residues 292-350) of TRAF6, were exclusively methylated in PIMT expressing cells, but not in PIMT depleted cells (Figure 5D). Interestingly, in addition to methylation, N350 deamination was also abundantly detected in both PIMT expressing and knockdown cells (Figure 5E), suggesting that the N350 deamination is naturally present regardless the levels of PIMT in cells. Since N350 is followed by a glycine, which is a typical “hot spot” for deamidation and subsequent *O*-methylation by PIMT (Reissner & Aswad, 2003), we speculate that deamination of N350 can be further methylated in PIMT expressing cells and this has been confirmed in our mass spectrometric analysis (Figure 5D and 5E).

D339 and N350 sites are found to be conserved among species (Figure 5F), reminding that these two sites might be critically for TRAF6 fuction. We generated two methylation-dead TRAF6 mutants (D339A and N350A) and examined the effects of these mutants on NF-κB activation by performing luciferase reporter assays. As shown in Figure 5G, overexpression of D339A and N350A mutants in EA.hy926 cells significantly elevated NF-κB transactivation to the similar extent as wild type TRAF6. Importantly, co-overexpression of PIMT markedly inhibited both wild-type TRAF6- and D339A mutant-induced NF-κB transactivation, but barely affected N350A-induced NF-κB transactivation, suggesting that methylation of TRAF6 at N350 is responsible for the PIMT-mediated inhibition of NF-κB activation. Furthermore, N350A mutation had no significant effect on TRAF6 oligomerization and auto-ubiquitination in the absence of PIMT, but markedly attenuated PIMT-mediated inhibition of TRAF6 oligomerization and auto-ubiquitination (Figure 5H and I). Together, these findings suggest that methylation of TRAF6 at N350 is critically involved in the regulation of TRAF6 function and NF-κB activation by PIMT.

### 6. PIMT impedes ICAM-1 expression and glycosylation in ECs

Activation of NF-κB has been implicated in the cytokine-induced expression of adhesion molecules, such as VCAM-1 and ICAM-1, which mediate the adhesion of monocytes to inflamed ECs (Pober & Sessa, 2007). Next, we determined whether PIMT affects cytokine-induced expression of VCAM-1 and ICAM-1 in ECs. As shown in Figure 6A, overexpression of PIMT markedly attenuated TNF-α-stimulated expression of ICAM-1 and VCAM-1 at the molecular weights of 110 kDa and 120 kDa, respectively, in a dose-dependent manner. Surprisingly, we observed that overexpression of PIMT1 also dose-dependently increased protein levels of ICAM-1 and VCAM-1 at lower molecular weights of 70 kDa and 90 kDa, which are speculated to be hypoglycosylated forms of these proteins. Similar results were observed in LPS- and IL-1β-treated HUVECs (Figure 6B) and HLMVECs (Supplementary Figure 3A), illustrating the broad relevance of our findings in endothelial biology. In attempt to determine whether PIMT impacts the global glycosylation pattern in ECs, we performed western blot using biotinylated lectin *Phaseolus vulgaris* leucoagglutinin (PHA-L) coupled with fluorescence labeled streptavidin staining of HUVEC lysates. No significant disparity was observed in Ad-LacZ- and Ad-PIMT-transduced ECs (Figure 6C). Additionally, we found PIMT knockdown by lentiviral shRNA efficiently increased ICAM-1 protein levels in both HUVECs and HLMVECs (Figure 6D). Since ICAM-1 is essentially involved in both adhesion and extravasation of leukocytes to endothelium during inflammation, it was chosen to further define the molecular mechanism of how ICAM-1 glycosylation was regulated by PIMT1. To this end, we performed co-IP to examine the interaction of PIMT1 with ICAM-1 in ECs. Our results showed that PIMT interacts only with hypoglycosylated ICAM-1 (∼70 kDa), but not with fully glycosylated form (∼110 kDa) (Figure 6E; Supplementary Figure 3B), suggesting that the interaction of PIMT with ICAM-1 occurs before ICAM-1 is full glycosylated and it potentially impairs the early processing of ICAM-1 glycosylation in ECs.

**Figure 6.**
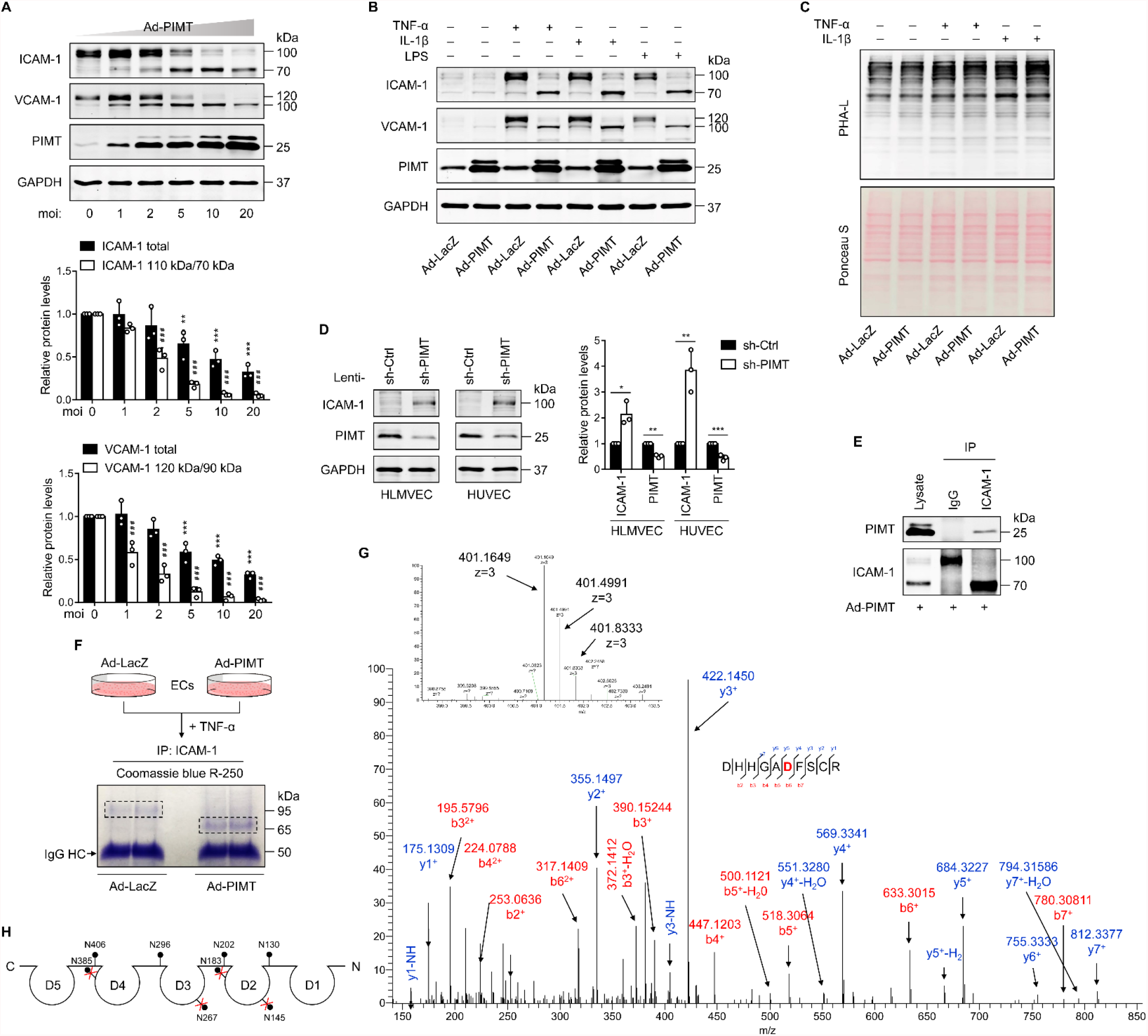
PIMT attenuates ICAM-1 expression and N-linked glycosylation in ECs. **A**, HUVECs were transduced with increasing dosages of Ad-PIMT for 48 hrs and then stimulated with TNF-α (10 ng/ml) for 12 hrs. Expression of ICAM-1 and VCAM-1 was determined by western blot. Total protein levels of ICAM-1 and VCAM-1 and the ratio of fully glycosylated bands to hypoglycosylated bands were quantitated at different mois (normalized to GAPDH), n = 3. \*\**P* < 0.01, \*\*\**P* < 0.001, compared with the corresponding ICAM-1 or VCAM1 expression at moi of 0. ^# # #^*P* < 0.001, compared with the corresponding the ratio of fully glycosylated bands to hypoglycosylated bands of ICAM-1 or VCAM-1 at moi of 0, two-way ANOVA coupled with Tukey’s post hoc test. **B**, HUVECs transduced with indicated ad-virus were exposed to TNF-α (10 ng/ml), IL-1β (20 ng/ml) or LPS (1 μg/ml) for 12 hrs, expression of ICAM-1 and VCAM-1 were detected by western blot. **C**, HUVECs were transduced with Ad-LacZ or Ad-PIMT (moi = 10) for 48 hrs and then treated with indicated cytokine and lysed for biotinylated PHA-L staining, followed by fluorescence labeled streptavidin incubation. Ponceau S staining reveals total loaded protein. **D**, HLMVECs and HUVECs were transfected with lentiviral control (sh-Ctrl) or PIMT (sh-PIMT) shRNA or for 72 hrs, expression of ICAM-1 was detected by western blot and quantitated by densitometric analysis. \**P* < 0.05, \*\**P* < 0.01, \*\*\**P* < 0.001, paired two-tailed t-test, n=3. **E**, HUVECs were transduced with Ad-PIMT (moi = 10) for 48 hrs then treated with TNF-α (10 ng/ml) for 12 hrs. Immunoprecipitation was performed using anti-ICAM-1 antibody, followed by western blot to detect the interaction of ICAM-1 with PIMT. **F**, Work flow of in gel sample preparation of ICAM-1 from ECs for LC-MS/MS analysis. Coomassie Blue R-250 staining showed immunoprecipitated ICAM-1 from HUVECs with indicated treatments. **G**, (Above) precursor ion peaks corresponding to 2 charged spectra, the m/z of the monoisotopic peak was exactly matched to the theoretical m/z (401.1649, z = 3). (Below) manually assigned MS/MS spectra of ICAM-1 peptide from PIMT overexpressed HUVECs. One representative spectrum is shown to exemplify the N-glycosite change. The Asn site was detected as Asp (shown in red font). **H**, Schematic demonstration of the N-glycosites of ICAM-1 (shown as lollipop patterns). Identified deamidated residuals were marked with red crosses.

ICAM-1 possesses a C-terminal cytoplasmic domain, a single transmembrane domain and an extracellular domain containing 8 N-glycosylation sites (Staunton *et al*., 1988). N-linked glycans are covalently attached to the amide side chain of Asn residues that resides in N-x-S/T consensus sequence (x represents any amino acid except proline). N-glycosylation impairment could induce major molecular weight shift (Mellquist *et al*, 1998). To determine whether PIMT affects the glycosylation of ICAM-1 at specific Asn sites, ICAM-1 was immunoprecipitated from the control and PIMT overexpressing ECs (Figure 6F) and then subjected to LC-MS/MS analysis to characterize the status of ICAM-1 N-glycosylation. Eight Asn residuals have been described as N-glyco sites in ICAM-1 in the previous studies (Casasnovas *et al*, 1998; Shimaoka *et al*, 2003; Yang *et al*, 2004), while in PIMT overexpression samples, four of which were identified to be switched to Asp residues, which are incapable of linking N-glycans (Figure 6G; Supplementary Figure 3C-E). Interestingly, all these sites were positioned in the antigen binding regions of ICAM-1 extracellular Ig domains (Figure 6H), suggesting that alteration of N-glycosylation at these sites by PIMT may change the interaction of ICAM-1 with its ligand. Taken together, these results demonstrate that PIMT prevents N-linked glycosylation of ICAM-1 presumably through enzymatic conversion of Asn to Asp at specific sites.

### 7. PIMT inhibits the function of ICAM-1 in endothelial activation

The canonical function of endothelial ICAM-1 and other adhesion molecules is involved in leukocyte arrest and extravasation (Springer, 1994). To determine whether PIMT affects ICAM-1-mediated function in ECs, we firstly examined the membrane location of ICAM-1 by flow cytometry and cell surface biotinylation assay. As shown in Figure 7A, TNF-α significantly increased cell surface expression of ICAM-1, which was markedly attenuated by ectopic expression of PIMT in ECs. Furthermore, cell surface biotinylation assay revealed that both the full-glycosylated and hypo-glycosylated forms of ICAM-1 were localized to the cell surface (Figure 7B), suggesting that glycosylation deficient ICAM-1 is able to escape the surveillance of endoplasmic reticulum (ER) quality control (ERQC) and localize to the cell membrane (Jørgensen *et al*, 2003). In addition, we found that PIMT overexpression markedly inhibited ICAM-1 membrane localization and its association with cytoskeletons, as determined by cellular fractionation and western blot. Notably, the hypo-glycosylated ICAM1 is totally absent in cytoskeletal fraction (Figure 7C). Next, we examined the effects of hypoglycosylation on ICAM-1 protein stability and found that the full-glycosylated ICAM-1 is stably preserved in HUVECs regardless of PIMT levels, while the degradation rate of hypo-glycosylated ICAM-1 is markedly increased, suggesting that PIMT impairs ICAM-1 glycosylation, which subsequently leads to increased degradation of ICAM-1 in ECs (Figure 7D). Monocyte adhesion to ECs is an important event in the initiation of proinflammatory response. Therefore, we examined the effect of PIMT on THP1 cell adhesion to the activated HUVECs. As shown In Figure 7E, TNF-α stimulation substantially increased THP1 cell adhesion to ECs, which was markedly inhibited by overexpression of PIMT. Cross-linking of ICAM-1 on the cell surface has been previously shown to cause intracellular signaling activation, which requires the interplay of ICAM-1 with cytoskeleton (Lawson *et al*, 1999). To determine whether PIMT affects ICAM-1-mediated cell signaling events in ECs, we ligated cell surface ICAM-1 using a monoclonal antibody in HUVECs, followed by cross-linking with a secondary antibody and examined the ICAM-1 inside-out signaling (Lawson & Wolf, 2009). As shown in Figure 7F, phosphorylation of ERK1/2 was efficiently induced by specific antibody ligation but not by control IgG, and this signal was suppressed by overexpression of PIMT. Pre-treatment with tunicamycin (TM), which completely blocks ICAM-1 glycosylation, also significantly attenuated the ICAM-1 crosslinking aroused signaling, further suggesting the role of glycosylation in ICAM-1-mediated signaling events. Together, these results suggest that PIMT functions as a negative regulator of the cytokine-induced inflammatory responses in vascular ECs, at least in part, through inhibiting both expression and glycosylation of adhesion molecules in vascular ECs.

**Figure 7.**
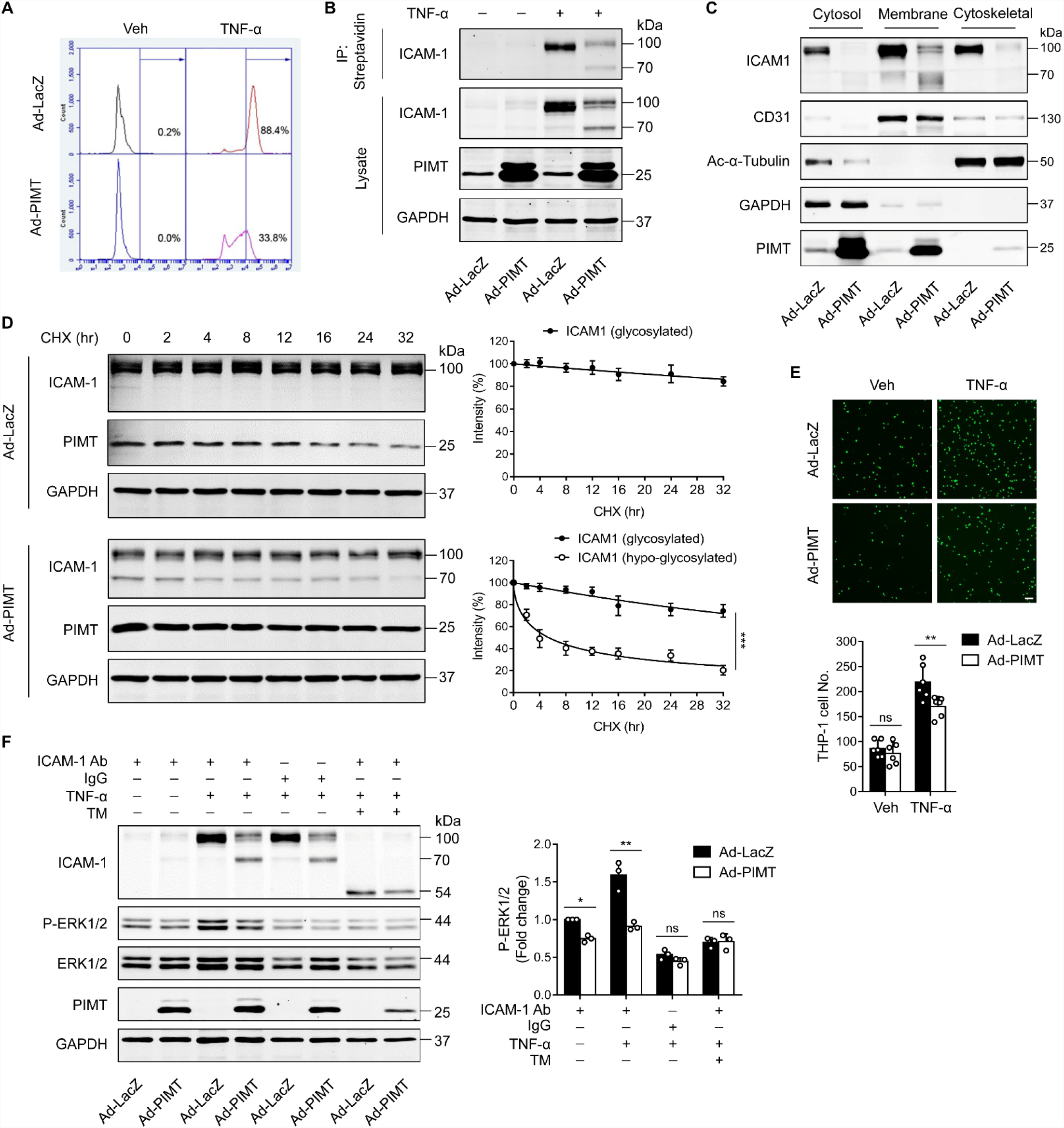
PIMT inhibits ICAM-1-mediated signaling events in ECs. **A**, HUVECs transduced with indicated adenovirus (moi = 10, 48 hrs) were stimulated with either vehicle or TNF-α (10 ng/ml) for 12 hrs. Expression of ICAM-1 on the surface of HUVECs was labeled by fluorescent antibody and determined by flow cytometry. **B**, HUVECs were transduced with Ad-LacZ or Ad-PIMT (moi = 10) for 48 hrs and then stimulated with TNF-α (10 ng/ml) for 12 hrs and labeled with biotin. Precipitation was performed using streptavidin agarose beads, followed by western blot to detect the cell surface ICAM-1. **C**, Western blot of ICAM1 in different cellular fractions. HUVECs were fractionated after indicated virus transduction (moi, 10) and TNF-α (10 ng/ml, 12hrs) incubation. GAPDH, CD31 and acetyl-α-tubulin were used as cytosol, membrane and cytoskeleton markers, respectively. **D**, HUVECs were transduced with Ad-LacZ or Ad-PIMT (moi = 5, 48 hrs) and stimulated with TNF-α (10 ng/ml) for 12 hrs to induce both ICAM-1 bands, then subjected to the treatment with 10 μM cycloheximide (CHX) for indicated intervals. The half-life of ICAM-1 was determined by western blot (n = 3), \*\*\**P* < 0.001, unpaired two-sided t test, **E**, HUVECs were stimulated with TNF-α (10 ng/ml, 12 hrs) and then incubated with calcein-labeled THP-1 cells for additional 1 hr. After washing, attached THP-1 cells were visualized and counted on an inverted fluorescent microscopy (n = 6), \*\**P* < 0.01, two-way ANOVA coupled with Tukey’s post hoc test. Bars, 50 μm. **F**, HUVECs pre-incubated with or without tunicamycin (TM, 2.5 μM) for 1 hr were exposed to indicated treatments and then subjected for ICAM-1 ligation using anti-ICAM-1 antibody followed by secondary antibody incubation, IgG were used as a negative cross-linking control. Phosphorylated and total ERK1/2 were determined by western blot and quantified by densitometric analysis (n = 3), \**P* < 0.05, \*\**P* < 0.01, two-way ANOVA coupled with Tukey’s post hoc test.

## Discussion

PIMT (encoded by Protein-L-Isoaspartate [D-Aspartate] *O*-Methyltransferase [*PCMT1*] gene) is a protein carboxyl methyltransferase which specifically recognizes structurally altered carboxylic acids stemmed from aspartyl residues and its role in monitoring protein homeostasis has been defined as “housekeeping” in a wide variety of cells and tissues (Desrosiers & Fanélus, 2011; Mishra & Mahawar, 2019; Reissner & Aswad, 2003). In this study, we identified a novel function of PIMT in suppressing endothelial activation and acute inflammatory responses. Specifically, we found that PIMT attenuates LPS-stimulated endothelial activation and acute lung injury by inhibiting NF-κB signaling through methylation of TRAF6 and inhibition of N-glycosylation of the cell surface adhesion molecule ICAM-1.

TRAF6 plays key roles in eliciting IL-1R/TLR mediated inflammatory signaling (Akira & Takeda, 2004; Chen, 2005; Deng *et al*., 2000). TRAF6 consists of an N-terminal RING domain, followed by four ZnF domains, a CC domain and a TRAF-C domain at the C-terminus (Chung *et al*., 2002). The RING domain, in association with ZnF domains, interacts with Ub-conjugating enzymes (E2) for Ub transfer (Yin *et al*, 2009). The TRAF-C domain is a binding scaffold mediating TRAF6 interaction with various effectors (Ye *et al*, 2002). Moreover, the CC domain has been shown to be indispensable for TRAF6 oligomerization and subsequent activation of TAK1 complex (Wang *et al*., 2006). Recently, the CC domain has been shown to enhance TRAF6 activation through interacting with E2 Ubc13 (Hu *et al*, 2017). However, little is known about how the function of TRAF6 CC domain is regulated under inflammatory conditions. Herein, we provide compelling evidence demonstrating that PIMT-mediated methylation of TRAF6 in the CC-domain represents a novel mechanism in regulating the TRAF6/NF-κB signaling in vascular ECs. Overexpression of PIMT markedly impeded TRAF6 oligomerization and auto-ubiquitination, which represent a key process in the initiation of LPS-induced NF-κB activation in ECs. Furthermore, we identified that the Asn residue 350 in the CC domain of TRAF6 is responsible for TRAF6 methylation by PIMT in ECs.

Deamidation is the first step of Asn to form PIMT recognizable intermediates, such as isoAsp, succinimide and methyl-isoAsp, during the protein repair process, and represents an important mechanism in regulating protein activity during development and/or aging of cells (Mishra & Mahawar, 2019; Reissner & Aswad, 2003) (Lee *et al*, 2012). Our data demonstrated that Asn substitution with Ala only impaired TRAF6 susceptibility to PIMT-mediated inhibition, but not the protein intrinsic functions, strongly indicating that PIMT catalyzed methylation, instead of repairing the deamidated end product, is the primary mechanism coordinating TRAF6 homeostasis. Interestingly, we found that the N350 residue in the CC domain is localized in a loop site adjacent to TRAF-C domain and exposed to the protein surface, which renders it more susceptible to deamidation (Supplementary Fig. 2) (Ye *et al*., 2002). Based on these findings, we extrapolated that the PIMT dependent N350 methylation serves as an intrinsic negative regulator of TRAF6 biological activity. It is worth noting that in this study we primarily focused on the functional interaction of PIMT1 with TRAF6 because TRAF6 is critically implicated in LPS-induced endothelial activation and acute lung injury (Chen, 2005; Martin & Wurfel, 2008). Our data also demonstrated a weak interaction between PIMT and TRAF2. Since a conservative deamidation susceptible Asn site is identified in the C terminal TRAF-C domain of TRAF2, whether PIMT1-mediatd *O*-methylation is involved in inhibiting TRAF2 activation warrants further investigation.

ECs are regarded as the sentinels for sensing invading pathogens in the circulation and triggering innate immune system activation, however, extravagant or hyperactive inflammatory response resulted from unrestrained EC activation can lead to detrimental effects on the host. Hence confining inflammatory reactions is imperative for governing the magnitude and duration of endothelial activation and immune responses. Indeed, several molecules, such as IκBs, deubiquitinase A20, transcriptional co-factor CITED2 and protein chaperone cyclophilin J, have been identified as critical negative regulators to inhibit inflammatory signals under various conditions (Boone *et al*, 2004; Lou *et al*, 2011; O’Dea & Hoffmann, 2010; Sheng *et al*, 2018). In the present study, we identified PIMT as a novel regulator in counteracting inflammatory responses through acting on TRAF6. We found that endotoxin treatment did not alter endothelial PIMT levels under both *in vitro* and *in vivo* conditions, but significantly augmented the interaction of PIMT and TRAF6 in different types of ECs, suggesting that the dynamic formation of PIMT/TRAF6 protein complex constitutes an important negative feedback mechanism in resolving LPS inflammation. Whether this enzyme possesses a role in the pathogenesis of other infectious diseases is under active investigation.

As a hallmark of endothelial activation, cell-surface expression of adhesion molecules mediates the interaction of circulating leukocytes with the endothelium in response to proinflammatory signals (Hunt & Jurd, 1998). Adhesion molecules, such as ICAM-1 and VCAM-1, are heavily N-glycosylated, and N-glycosylation defects have been shown to impair the expression and membrane localization of adhesion molecules and subsequent intracellular, cell–cell and cell–matrix recognition events in inflammatory responses (Chen *et al*., 2014b; Scott & Patel, 2013). N-glycan only attaches to Asn that presents in an N-x-S/T consensus sequence. At this time, little is known about how N-glycosylation of adhesion molecules is regulated under inflammatory conditions in ECs. Here in this work, we uncovered an unexpected role of PIMT in modulating N-glycosylation of adhesion molecules ICAM-1 and VCAM1. We found that PIMT specially inhibits the N-glycosylation of ICAM-1 and VCAM-1 without altering the global endothelial glycocalyx. To further investigate molecular mechanism underlying inhibition of adhesion molecule N-glycosylation by PIMT, we chose to determine the structure of the N-glycans of ICAM-1 using mass spectrometry. We found that PIMT inhibits ICAM-1 N-glycosylation at the specific Asn sites (N145, N183, N267 and N385). Our mass spectrometric data show that in the presence of PIMT, Asn residues at these sites are switched to Asp residues, thus preventing N-glycosylation. In addition to decreasing ICAM-1 protein stability as shown in this study, we expect that the deficiency of ICAM-1 N-glycosylation at these specific Asn sites may directly impact the function of ICAM-1, despite our data show that hypo-glycosylated form of ICAM-1 is able to localize to the cell membrane. As shown in Figure 6H, the N145 and N183 sites are localized in the hydrophilic BC and EF loops of ICAM-1 domain 2 (D2) upon structural dissection (Casasnovas *et al*., 1998). Based on the fact that ICAM-1 D2 sustains the structural and LFA1 binding integrity of D1 (Stanley *et al*, 2000), we speculate that PIMT-induced “site mutation” at these sites may impair the binding capacity of ICAM-1 to its ligand, and this hypothesis was partially testified in our ICAM-1 antibody cross-linking experiments. Furthermore, lack of ICAM-1 glycosylation at N267 in the D3 domain may also affect its interaction with macrophage-1 antigen (Mac-1) (Diamond *et al*, 1993). In this regard, our study provides novel insights into how PIMT-mediated crosstalk between O-methylation and N-glycosylation regulates the functional interaction of ECs and immune cells in vascular inflammation. Elucidation of molecular mechanisms underlying PIMT-induced hypo-glycosylation of VCAM-1 in ECs is ongoing.

In summary, our results have identified novel mechanisms in controlling EC activation by PIMT in vascular inflammation. We demonstrate that PIMT-catalyzed methylation directly restricts TRAF6 activation to maintain TRAF6 in the default static state. Moreover, PIMT provides a “double check” mechanism to regulate ICAM-1 membrane expression and intracellular signaling at both transcriptionally and post-translational levels to avoid endothelial activation (Figure 8). Our results suggest that targeted activation of PIMT may provide a novel therapeutic strategy for the treatment of inflammatory vascular disorders.

**Figure 8.**
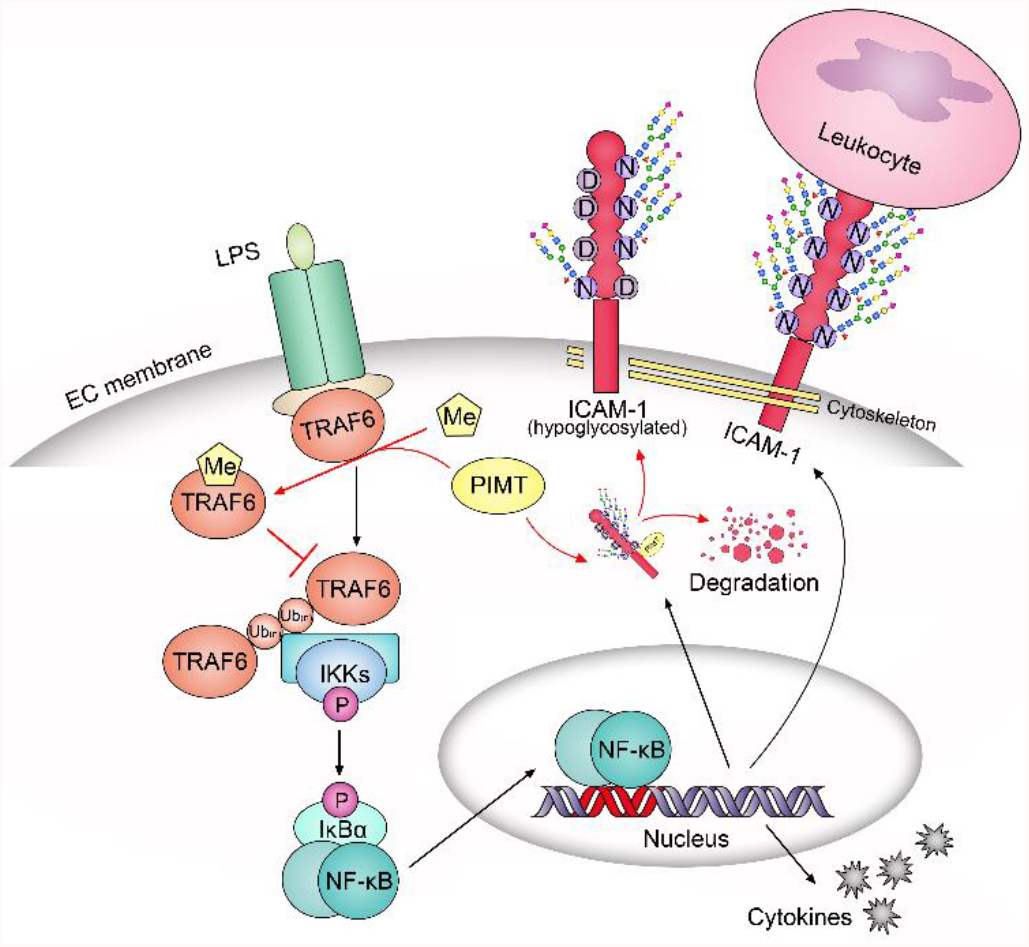
Schematic diagram of suppressing LPS-stimulated endothelial activation by PIMT. PIMT prevents LPS-induced EC activation through methylation of TRAF6 and inhibition of ICAM-1 expression and glycosylation, hence serving as a negative feedback approach to confine extravagant endothelial inflammatory response.

## Materials and Methods

### Mice

PIMT heterozygous knockout mice (B6;129S4-Pcmt1^tm1Scl^/J, #023343) were purchased from The Jackson Laboratory and bred with the WT counterparts (C57BL/6J, #000664, JAX) to maintain live colonies. Genotyping PCR were performed using genomic DNA by tail snipping. The WT (5’-AGTGGCAGCGACGGCAGTAACAGCGG-3’ and 5’-ACCCTCTTCCCATCCACATCGCCGAG-3’) and knockout primer sets (5’-CGCATCGAGCGAGCACGTACTCGG-3’ and 5’-GCACGAGGAAGCGGTCAGCCCATTC-3’) targeting PCMT1 exon 1 or neomycin cassette were described elsewhere before(Dai *et al*, 2013). Mixed male and female mice at ages of 8-10 weeks were used in the study to induce acute lung injury. Protocols involving animal care and use were approved by the Institutional Animal Care and Use Committee at Thomas Jefferson University before initiation of any studies.

### Cell culture

Human umbilical vein endothelial cells (HUVEC, Gibco, C0155C) and human lung microvascular endothelial cells (HLMVEC, Lonza, CC-2527) were cultured in complete endothelial cell medium (ScienCell, 1001) and used within 3 passages. HEK-293T (ATCC, CRL-3216), EA.hy926 (ATCC, CRL-2922) and 293AD cells (Cell Biolabs, AD-100) were maintained in DMEM medium (Corning, 10013CV), THP-1 cells (ATCC, TIB-202) were cultures in RPMI-1640 medium (Corning, 10-040-CM), all supplemented with fetal bovine serum (FBS) (Gibco, 1082147) and penicillin/streptomycin (Corning, 30-002-CI).

### Reagents and antibodies

Anti-PCMT1 (A6684), anti-TRAF6 (A0973) anti-ICAM1 (A5597) and anti-HA (AE036) antibodies were from ABclonal Technology. Anti-PECAM1 (550274) antibody was from BD Pharmingen. Anti-Flag (F3165), anti-HA (H9658), anti-c-Myc (C3956) antibodies, Tunicamycin (11089-65-9) and Cycloheximide (C7698) were from Sigma-Aldrich. Anti-GAPDH (10494-1-AP), anti-alpha Tubulin (11224-1-AP), anti-TRAF6 (66498-1-Ig) and anti-VEGFR2 (26415-1-AP) antibodies were from Proteintech Group. Anti-p65 (D14E12), anti-p105/p50 (D4P4D), anti-Phospho-IKKα/β (#2697), anti-IKKβ (D30C6), anti-Phospho-IκBα (14D4), anti-IκBα (#9242), anti-Phospho-p44/42 MAPK (Erk1/2) (D13.14.4E), anti-p44/42 MAPK (Erk1/2) (#4695) and anti-Acetyl-α-Tubulin (Lys40) (#5335) antibodies were from Cell Signaling Technology. Anti-Lamin A/C (sc-20681), anti-ICAM1 (sc-7891), anti-VCAM1 (sc-13160), anti-PCMT1 (sc-100977) antibodies and LPS (sc-3535) were from Santa Cruz Biotechnology. IRDye 680RD Donkey anti-Rabbit (926-68073), 800CW Goat anti-Mouse (926-32210) antibodies and 680RD Streptavidin (926-68079) were from LI-COR Bioscience. Alexa Flour 488 donkey anti-mouse (A21202), 488 donkey anti-rat (A21208), 555 goat anti-rabbit (A21428), 647 goat anti-rabbit (A32733) antibodies, CellTracker Green CMFDA Dye (C2925) and DAPI (4’,6-diamidino-2-phenylindole) (D1306) were from Invitrogen. EZ-Link Sulfo-NHS-LC-LC-Biotin (#21135) and Streptavidin Agarose (#20357) were from Pierce. Biotinylated Phaseolus Vulgaris Leucoagglutinin (PHA-L, B-1115) was from Vector Laboratories. Recombinant Human TNF-α (300-01A) and IL-1β (200-01B) were from PeproTech.

### Plasmids

pFLAG-CMV-2-PIMT construct was described previously (Yan *et al*., 2013). Myc-PIMT was generated by cloning the PIMT fragment into pCS2-6×Myc plasmid. pcDNA3-FLAG-TRAF6 (#66929), pcDNA3-FLAG-TRAF2 (#66931), pcDNA-FLAG-IKKβ (#23298), FLAG-IKKβ-S177E/S181E (#11105) and HA-Ubiquitin (#18712) were acquired from Addgene. HA-TRAF6 was generated by cloning the TRAF6 fragment into pcDNA3-HA plasmid, and HA or Myc tagged TRAF6 truncations were made from pcDNA3-FLAG-TRAF6 or pcDNA3-HA-TRAF6 for direct transfection. PIMT and TRAF6 mutants were made using the QuikChange II Site-Directed Mutagenesis Kit (Agilent, #200523). All original constructs in this study were verified by DNA sequencing.

### Bronchoalveolar lavage

Bronchoalveolar lavage (BAL) was performed by cannulating the trachea with a blunt 22-gauge needle and lavaging with 1 ml PBS as described before (Zhu *et al*, 2019). Total cell counts were determined using a TC20 automated cell counter (Bio-Rad). After centrifugation, BAL fluid supernatant was subjected to protein concentration measurement by BCA assay (Pierce, #23225). BAL fluid neutrophils were re-suspended in PBS and analyzed after cyto-centrifugation onto glass slides and staining with Kwik-Diff solutions (Thermo Scientific, #9990701).

### Lung histology

Lung histology was performed on paraformaldehyde fixed tissues embedded in paraffin wax. Sections (5 μm) were placed on positively charged glass slides, and then deparaffinized for hematoxylin and eosin (H&E) staining. Tissues were visualized with a light microscope (Nikon).

### Enzyme-linked immunosorbent assay (ELISA)

Mouse BAL fluids were collected for ELISA measurement of TNFAα (DY-410), IL-6 (DY-406) and Ccl2 (DY-479) according to the manufacturer’s manual (R&D Systems). Briefly, BAL fluids and standards were added in various dilutions to the plate coated with capture antibody, and then labeled with biotinylated detection antibody. After incubation with streptavidin-peroxidase and substrate, protein concentration was determined by absorbance at 450 nm in a plate reader (Biotek Instrument).

### Adenovirus construction and purification

Adenoviruses were generated using RAPAd adenoviral expression systems (Cell Biolabs) according to the manual. Briefly, pacAd5 CMVK-NpA shuttle vector containing gene of interest and pacAd5 9.2-100 backbone vector were linearized using PacI digestion, and then co-transfected to 293AD cells. Crude viral lysates were harvested from the adenovirus-containing cells by three freeze/thaw cycles.

The viral supernatants were propagated in 293AD cells to amplify, and purified through CsCl density-gradient ultracentrifugation, followed by dialysis into 20 mM Tris buffer and supplement with 10% glycerol for stocking. Viral titers were determined using the Adeno-X Rapid Titer Kit (Takara Bio, 632250).

### Transient Transfection

DNA was transfected to host cells using polyethylenimine (PEI). In general, cells were seeded to be 80% confluent at transfection. Plasmid DNA was mixed with PEI (1 mg/ml) at 1:3 in Opti-MEM medium (Gibco, 11058021) and added to cells after short incubation. Medium was changed after 6 hr, and cells were harvested after 48 hr incubation.

siRNA Universal Negative Control (SIC001) and si-PCMT1 (SASI_Hs01_00229688) were from Sigma-Aldrich and transfected using Lipofectamine RNAiMAX Transfection Reagent (Invitrogen, 13778150) according to the recommendations of the manufacturer. Transfected cells were verified after 72 hr.

### Lentivirus infection

Stable cell lines were made by lentivirus transduction, as previously described (Chen *et al*, 2014a) with slight modifications. pLKO-PCMT1 construct (TRCN0000036403) and Non-Target shRNA Control (SHC016) were from Sigma-Aldrich. Lentiviral pLKO constructs were transfected with packaging and envelope plasmids to HEK293T cells. Viral supernatant was harvested 48 hr and 72 hr post-transfection, and filtered through a 0.45 μm filter before adding to recipient cells, and supplemented with 10 μg/ml polybrene. The infected cells were selected with puromycin (1 μg/ml) for ∼1 week.

### Luciferase reporter assay

NF-κB firefly luciferase reporter vectors (pNF-κB-TA-Luc) and Renilla luciferase control reporter vectors (pRL-TK), together with indicated expression plasmids were transfected into cells pre-seeded in 24-well plates. Luciferase level was measured as described previously by a luminescence microplate reader (BioTek). Basically, cells were lysed 48 hr later and firefly luciferase was detected using 2×GoldBio Luciferase Assay Buffer containing 200 mM Tris (pH 7.8), 10 mM MgCl_2_, 0.5 mM Coenzyme A (CoA), 0.3 mM ATP, 0.3 mg/ml Luciferin (Gold Biotechnology, #LUCK). Renilla luciferase was assessed using 2 μM coelenterazine in DPBS. For LPS or IL-1β stimulated reporter activity, cells were incubated for 10 hr with designated treatments and lysed.

### Western blot

Cells were lysed in RIPA buffer containing 50 mM Tris (pH 8.0), 0.5 mM EDTA, 150 mM NaCl, 1% NP-40, 1% sodium dodecyl sulfate (SDS), supplemented with protease and phosphatase inhibitors. The lysates were agitated, and supernatants were denatured and subjected to SDS-PAGE. After transferring, the nitrocellulose membranes were blocked with 5% BSA at room temperature for 1 hr. Primary antibodies were incubated at 4°C overnight. After washing, the membrane was incubated with IRDye secondary antibody conjugates at room temperature for 1 hr. The membrane was then washed and imaged by the Odyssey infrared imaging system (LI-COR, 9120).

### Immunofluorescence

Cells plated on 8-well Chamber Slides (Thermo Scientific, 154534) with indicated treatments were fixed with 4% formaldehyde and permeabilized with 0.2% Triton-X 100 at room temperature. After blocking with 5% BSA for 1 hr, slides were incubated with primary antibodies overnight at 4°C, followed by exposure to Alexa Fluor-conjugated secondary antibodies for 1 hr at room temperature. After staining with DAPI (1 μg/ml), the wells were removed and slides were mounted with 50% glycerol and imaged with a fluorescent confocal microscope (Nikon). For staining of lung frozen sections, mouse lung were inflated with OCT before snap freezing and sectioned a thickness of 5 μm, followed by immunostaining.

### Immunoprecipitation

Cells were lysed in IP buffer containing 20 mM Tris (pH 8.0), 137 mM NaCl, 2 mM EDTA, 1% NP-40, and 10% glycerol, supplemented with protease and phosphatase inhibitors. For endogenous IP, the supernatants were pre-cleaned with IgG and Dynabeads (Thermo Fisher Scientific), then incubated with antibodies at 4°C overnight with rotation, indicated beads were added for additional 2 hr on the next day. For exogenous IP, the supernatants were incubated with Anti-FLAG M2 Magnetic Beads (Sigma-Aldrich) and rotated at 4°C overnight. The IP was washed 3 times with lysis buffer, collected by magnet, and then boiled with SDS reducing loading dye. Samples were analyzed by western blot.

### Subcellular fractionation

Nucleus and cytosol isolation was performed with the NE-PER Nuclear and Cytoplasmic Extraction Reagents (Thermo Scientific, #78835) according to the manufacturer instructions. The nuclear extracts here were used for EMSA assay. Extraction of cell membrane and cytoskeleton fractions was carried out using the Subcellular Protein Fractionation Kit for Cultured Cells (Thermo Scientific, #78840) in line with the manual.

### Electrophoretic mobility shift assay (EMSA)

EMSA was performed with IRDye 700 NFĸB Consensus Oligonucleotide (LI-COR, 829-07924) according to the manufacturer instructions with slight modifications. Briefly, 10 μg nuclear extracts prepared from HUVECs were incubated with 40 fmol labeled NF-ĸB oligonucleotides at room temperature for 30 min, and DNA-protein complexes were separated in 4% native polyacrylamide gel containing 50 mM Tris (pH 7.5), 0.38 M glycine and 2 mM EDTA, the shift was detected by the Odyssey infrared imaging system. The binding specificity was determined using 50 fold unlabeled DNA duplex or anti-p65 antibody.

### Flow cytometry

Adenovirus transduced HUVECs were treated with TNFα for 12 hr to induce ICAM-1 and VCAM-1 expression. HUVECs were then trypsinized and resuspended in fluorescence-activated cytometry sorting (FACS) buffer containing 10% FBS and 1% sodium azide in PBS, followed by incubation with indicated primary antibodies for 1 hr on ice. After washing, cells were incubated with Alexa fluor conjugated fluorescent secondary antibodies for 30 minutes on ice in the dark. Then samples were washed again and resuspended in FACS buffer. For each sample, 1.5×10^4^ cells were analyzed by a BD Accuri C6 flow cytometer (BD Bioscience).

### Monocyte Adhesion Assay

HUVECs were seeded in 24-well plates pre-coated with 0.1% gelatin. After virus induction and stimulation with inflammatory cytokines, 2.5×10^5^ THP-1 cells labeled with calcein CellTracker Green CMFDA Dye (Invitrogen) were added to each well in serum free media and incubated for 30 min at 37°C. After washing, the number of attached THP-1 cells was counted on the EVOS FL Auto Imaging System (Invitrogen).

### Quantitative real-time RT-PCR

Quantitative real-time RT-PCR (qRT-PCR) was performed as described previously. Total RNA was extracted from cells with TRIzol reagent (Invitrogen), and cDNA was synthesized using the High-Capacity cDNA Reverse Transcription Kit (Applied Biosystems). RT-PCR was performed using the PowerUp SYBR Green Master Mix (Applied Biosystems) on a CFX Connect Real-Time PCR Detection System (Bio-Rad), and results were calculated by the comparative cycling threshold (Ct) quantification method. The gene encoding GAPDH was served as an internal control for the total amount of cDNA. The primer sequences used were described as follows: Human ICAM1, 5′-CTTCGATCCCAAGGTTTCCAA-3′ (F) and 5′- TCGACACAAAGGATTTCGTAAGG-3′ (R); VCAM1, 5′-GGGAAGATGGTCGTGATCCTT-3′ (F) and 5′- TCTGGGGTGGTCTCGATTTTA-3′ (R); TNFα, 5′- CCTCTCTCTAATCAGCCCTCTG-3′ (F) and 5′- GAGGACCTGGGAGTAGATGAG-3′ (R); IL-1β, 5′-ATGATGGCTTATTACAGTGGCAA-3′ (F) and 5′- GTCGGAGATTCGTAGCTGGA-3′ (R); IL-6, 5′-ATGAGGAGACTTGCCTGGTGAA-3′ (F) and 5′- AACAATCTGAGGTGCCCATGCTAC-3′ (R); Ccl-2, 5′-CAGCCAGATGCAATCAATGCC-3′ (F) and 5′- TGGAATCCTGAACCCACTTCT-3′ (R); GAPDH, 5′-ACAACTTTGGTATCGTGGAAGG-3′ (F) and 5′- GCCATCACGCCACAGTTTC-3′ (R); mouse ICAM1, 5’-GTGATGCTCAGGTATCCATCCA-3’ (F) and 5’-CACAGTTCTCAAAGCACAGCG-3’ (R); TNFα, 5’- CCCTCACACTCAGATCATCTTCT-3’ (F) and 5’- GCTACGACGTGGGCTACAG-3’ (R); IL-1β, 5′- GCAACTGTTCCTGAACTCAACT-3′ (F) and 5′- ATCTTTTGGGGTCCGTCAACT-3′ (R); IL-6, 5′- TAGTCCTTCCTACCCCAATTTCC-3′ (F) and 5′- TTGGTCCTTAGCCACTCCTTC-3′ (R); Ccl-2, 5′- TTAAAAACCTGGATCGGAACCAA-3′ (F) and 5′- GCATTAGCTTCAGATTTACGGGT-3′ (R); GAPDH, 5′- AGGTCGGTGTGAACGGATTTG-3′ (F) and 5′- TGTAGACCATGTAGTTGAGGTCA-3′ (R).

### Ubiquitination assay

Detection of protein ubiquitination in cultured cells was conducted as described previously (Choo & Zhang, 2009). Basically, cells were lysed with 2% SDS, 150 mM NaCl, 10 mM Tris (pH 8.0) supplemented with protease inhibitors. The lysates were boiled and sonicated, and diluted with 150 mM NaCl, 10 mM Tris (pH 8.0), 2 mM EDTA, and 1% Triton. After incubation at 4°C with rotation, the supernatants were collected for anti-Flag immunoprecipitation.

### Protein sample preparation and nano liquid chromatography coupled to mass spectrometry (nano LC-MS/MS)

Gel band was destained with 100 mM ammonium bicarbonate/acetonitrile. The band was reduced in 10 mM dithiothreitol/100 mM Ammonium bicarbonate and alkylated with 100 mM iodoacetamide in 100 mM ammonium bicarbonate. Proteins in the gel band were then digested with trypsin overnight and collected in supernatants. Additional gel peptides were extracted by 50% acetonitrile and 1% TFA. The supernatants were combined and dried, followed by reconstitution of 0.1% formic acid for mass spectrometry analysis.

Desalted peptides were analyzed on a Q-Exactive HF attached to an Ulimate 300 nano UPLC system (Thermo Scientific). Peptides were eluted with a 25 min gradient from 2% to 32% ACN and to 98% ACN over 5 min in 0.1% formic acid. Data dependent acquisition mode with a dynamic exclusion of 45 second was enabled. One full MS scan was collected with scan range of 350 to 1200 m/z, resolution of 70 K, maximum injection time of 50 ms and AGC of 1e6. Then, a series of MS2 scans were acquired for the most abundant ions from the MS1 scan (top 15). An isolation window of 1.4m/z was used with quadruple isolation mode. Ions were fragmented using higher-energy collisional dissociation (HCD) with a collision energy of 28%.

Proteome Discoverer 2.3 (Thermo Scientific) was used to process the raw spectra. Database search criteria were as follows: (taxonomy Homo Sapiens) carboxyamidomethylated (+57 Da) at cysteine residues for fixed modifications; Oxidized at methionine (+16 Da) residues; Phosphorylation (+79.9 Da) at serine, threonine, and tyrosine residues; Deamidation (+0.98 Da) and methylation (+14 Da) at asparagine/aspartate for variable modifications, two maximum allowed missed cleavage, 10 ppm MS tolerance, a 0.02-Da MS/MS tolerance. Only peptides resulting from trypsin digestion were considered. The target-decoy approach was used to filter the search results, in which the false discovery rate was less than 1% at the peptide and protein level.

### Statistical analysis

Data were presented as mean ± SD and plotted using GraphPad Prism version 7.00 software. Two-tailed Student’s t-test was applied for comparison between two groups. Two-way ANOVA coupled with Tukey’s post hoc test was applied for two independent variables. Significance was considered when the P value was less than 0.05.

## Data availability

This study includes no data deposited in external repositories.

## Acknowledgments

This work was supported by grants from National Institute of Health (RO1HL103869, RO1GM123047, RO1HL159168, RO1HL152703), and American Heart Association Established Investigator Award (16EIA27710023).

## Author contributions

C.Z. performed and interpreted experiments and wrote the manuscript. B.Y. performed the experiments. Z.G., K.K., W.L., X.S., L.W., X.Y, R.S interpreted experiments and revised the manuscript. J.S. conceived and supervised the study and revised the manuscript.

## Conflict of interest

The authors declare that they have no conflict of interest.

